# *leptin b* and its regeneration enhancer illustrate the regenerative features of zebrafish hearts

**DOI:** 10.1101/2022.07.14.500053

**Authors:** Kwangdeok Shin, Ian J. Begeman, Jingli Cao, Junsu Kang

## Abstract

Zebrafish possess a remarkable regenerative capacity, which is mediated by the induction of various genes upon injury. Injury-dependent transcription is governed by the tissue regeneration enhancer elements (TREEs). Here, we utilized *leptin b* (*lepb*), an injury-specific factor, and its TREE to dissect heterogeneity of non-cardiomyocytes (CMs) in regenerating zebrafish hearts. Our single-cell RNA sequencing (scRNA-seq) analysis demonstrated that the endothelium/endocardium(EC) is activated to induce distinct subpopulations upon cardiac injury. We demonstrated that *lepb* can be utilized as a regeneration-specific marker to subset injury-activated ECs. *lepb*^+^ ECs robustly induce pro-regenerative factors, implicating *lepb*^+^ ECs as a signaling center to interact with other cardiac cells. Our scRNA-seq analysis identified that *lepb* is also produced by specific subpopulation of epicardium (Epi) and epicardium-derived cells (EPDCs). To determine *lepb* labels injury-emerging non-CM cells, we tested the activity of *lepb*-linked regeneration enhancer (*LEN*) with chromatin accessibility profiles and transgenic lines. While non-detectable in uninjured hearts, *LEN* directs EC and Epi/EPDC expression upon injury. The endogenous *LEN* activity was assessed using *LEN* deletion lines, demonstrating that *LEN* deletion abolished injury-dependent expression of *lepb*, but not other nearby genes. Our integrative analyses identify regeneration-emerging cell types and factors, leading to the discovery of regenerative features of hearts.

## INTRODUCTION

Adult mammals poorly regenerate damaged hearts, resulting in high morbidity and mortality from cardiac diseases. In contrast, zebrafish possess a remarkable ability to regenerate injured hearts. Heart regeneration studies across animal species identified multiple regeneration-driving genes and signaling pathways, which are highly conserved between mammals and zebrafish ^1^. These results suggest that a key difference between mammals and zebrafish is not the presence or absence of regeneration-driving genes but in the mechanisms controlling expression of these genes after injury ^2–4^. Dissecting the regulatory mechanism governing zebrafish heart regeneration will provide insights into understanding the molecular basis of cardiac regeneration.

Heart regeneration is a complex process, in which cardiomyocytes (CMs) and non-CMs cooperatively play roles to trigger regenerative programs. As CMs are muscular cells that ensure cardiac functions to circulate blood throughout the body, an essential event for heart regeneration is to activate CM proliferation. Non-CMs, such as endocardium, epicardium (Epi), and immune cells, are also crucial cardiac tissues that respond to cardiac injury to initiate regenerative process. Endocardium and Epi in zebrafish are rapidly activated within several hours and one day post-injury, respectively, contributing to heart regeneration in multiple aspects^5, 6^. For instance, cardiac injury-activated endocardium and Epi produces paracrine factors to stimulate CM proliferation^5, 7–9^ and extracellular matrix proteins (ECMs) to construct the regenerative niche/environment^10, 11^. Activated endocardium and Epi are thought to be heterogenous^12–15^, but molecular identity representing diverse subgroups of these cardiac tissues in regenerating hearts is relatively unexplored.

Tissue regeneration enhancer elements (TREEs) are key regulatory elements that relay injury signals to direct gene expression ^3, 16–20^. We previously identified the first cardiac TREE in zebrafish, the *leptin b* (*lepb*)-linked regeneration enhancer (*LEN*) ^17^*. LEN* and *lepb*, the target gene of *LEN*, are not active during development but are robustly activated upon injury^21^, highlighting their regeneration-specific characteristics. Here, we dissect *lepb-* expressing cell populations at the single-cell level to infer dynamic changes of non-CM populations in injured hearts. Further studies of epigenome profiles, enhancer assays, and *LEN* deletion mutant analysis determine the regeneration-dependent specificity of *LEN* in non-CMs. Overall, our comprehensive analyses of multiple transcriptomic and epigenomic profiles of injured hearts identify novel molecular and cellular targets for heart regeneration and provide insights into the regeneration-specific features of the hearts.

## RESULTS

### scRNA-seq analysis combined with *lepb*, a regeneration-specific marker, enhances classification of heterogenous non-CM populations into subgroups

Injury-induced genes can be utilized for representing cell types emerging upon injury. Our previous work demonstrated that *lepb* exhibits regeneration-specific expression in endocardial cells^21^, indicating the potential advantage of *lepb* as a regeneration marker. To investigate transcriptomic changes at the single cell level in adult hearts, we analyzed available single cell sequencing (scRNA-seq) datasets generated with ventricular cells sorted for either *runx1P2:Citrine* or *kdrl:mCherry*^22^*. runx1P2:Citrine* expressed in a wide range of cells in injured hearts, including CMs, Epi, endocardium, and blood cells and *kdrl:mCherry* labelled endothelium/endocardium in the hearts^22^. Datasets obtained from wild-type uninjured and 3 days post-cryo-injury (dpi) hearts were used for unsupervised clustering, identifying 23 different clusters (**Fig. 1A, B and S1A, B**). To determine cell types composing these clusters, we annotated each cluster with known marker genes (**Fig. 1A, C and S1A, C**): CMs with *tnnt2a* and *myl7*^23, 24^; epicardial cells (Epi) and cardiac fibroblasts (cFB) with *tcf21, fn1b, col1a1a, tagln,* and *col5a1*^12, 25–27^; endocardial/endothelial cells (ECs) with *cdh5, kdrl,* and *flt1*^28, 29^; coronary endothelial cells (cEC) composing blood vessels in hearts with *cdh5, kdrl, flt1,* and *aplnra*^12^; mesenchyme-like cells (Mes) with *mgp, angptl7,* and *rspo1*^30, 31^; thrombocytes (throm) with *itga2b* and *gp1bb*^32, 33^; neutrophils (Neu) with *mpx* and *lyz*^34, 35^; macrophages (MC) with *mfap4, c1qa,* and *mpeg1.1*^36–38^; leukocytes (leu) with *mhc2a, coro1a,* and *cxcr4b* ^39–41^(**Table 1**).

**Figure 1.**
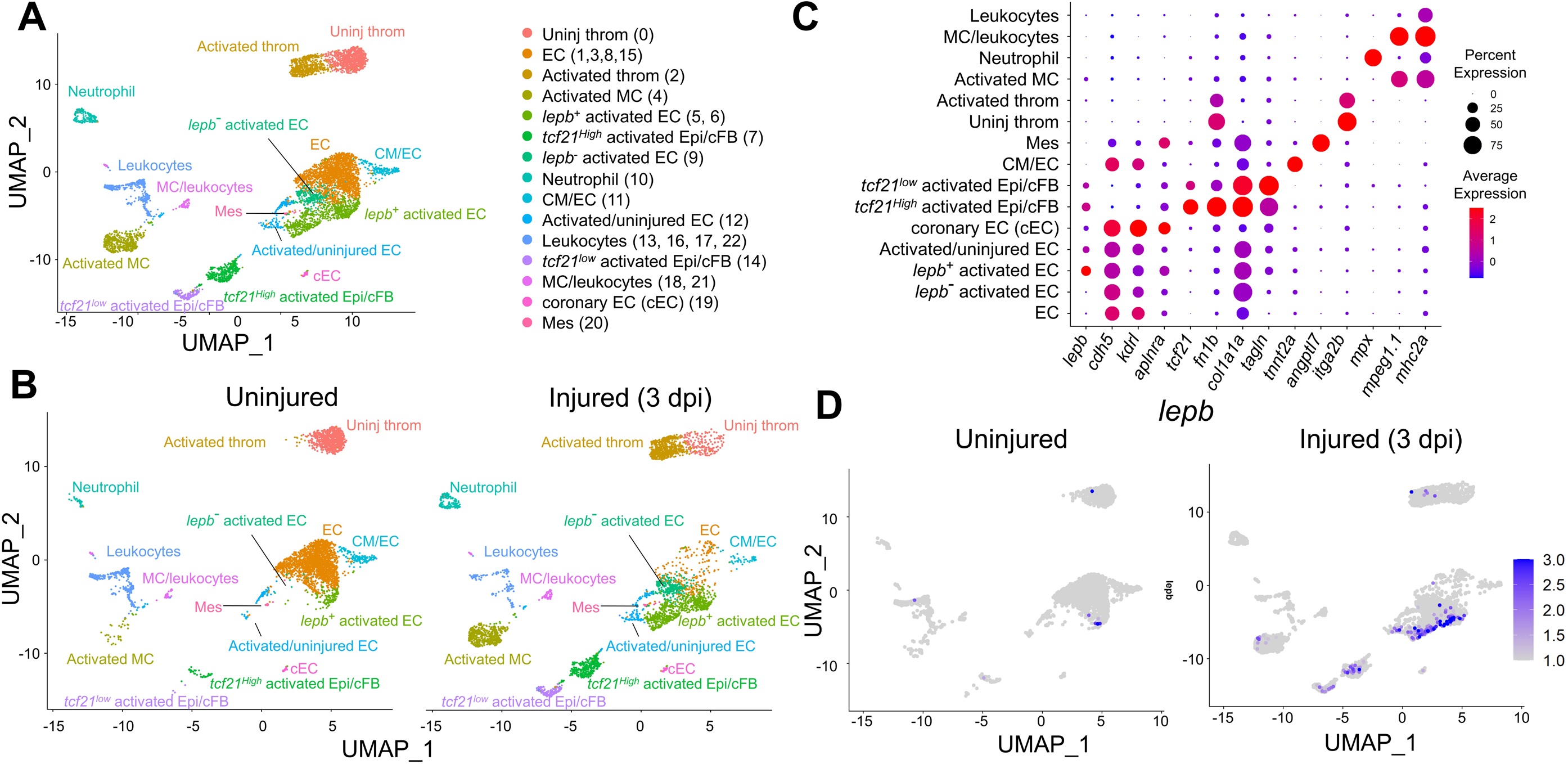
Heterogenous cell clusters in injured hearts of adult zebrafish. (A) Clustering assignments of *runx1P2:Citrine* or *kdrl:mCherry* expressing cells collected from uninjured and injured hearts. Uniform Manifold Approximation and Projection (UMAP) axes were calculated by unsupervised clustering method. throm, thrombocyte. EC, endocardial/endothelial cells. Epi, epicardial cells. cFB, cardiac fibroblasts. Mes, mesenchyme-like cells. MC, macrophages. (B) Clustering assignments for uninjured and injured hearts. dpi, days post-injury. (C) Differential expression of the key marker genes to identify cell types shown as a dot plot. (D) Injury-dependent expression of *lepb* depicted by UMAP plot.

**Table 1.** Cell number of injured and uninjured clusters identified by scRNA-seq analysis.

| Cluster | Injured (3dpi) | Uninjured | Total cell number | Marker genes |
| --- | --- | --- | --- | --- |
| EC (uninjured) | 225 | 1940 | 2165 | *cdh5, kdrl,flt1* |
| *lepb<sup>-</sup>* activated EC | 315 | 8 | 323 | *cdh5, kdrl,flt1* |
| *lepb<sup>+</sup>* activated EC | 740 | 128 | 868 | *cdh5, kdrl,flt1,lepb* |
| Activated/uninjured EC | 119 | 69 | 188 | *cdh5, kdrl,flt1* |
| coronary EC (cEC) | 41 | 19 | 60 | *cdh5, kdrl,flt1,aplnra* |
| *tcf21<sup>low</sup>* activated Epi/cFB | 177 | 4 | 181 | *tcf21,fn1b,col1a1a,tagln,col5a1* |
| *tcf21<sup>High</sup>* activated Epi/cFB | 381 | 44 | 425 | *tcf21,fn1b,col1a1a,tagln,col5a1* |
| CM/EC | 57 | 149 | 206 | *tnnt2a and myl7* |
| Mes | 24 | 10 | 34 | *mgp, angptl7,rspo1* |
| Uninjured thromcyte | 180 | 792 | 972 | *itga2b, gp1bb* |
| Activated thromcyte | 636 | 10 | 646 | *itga2b, gp1bb* |
| Activated Macrophage | 540 | 32 | 572 | *mfap4, c1qa, mpeg1.1* |
| Neutrophil | 176 | 41 | 217 | *mpx, lyz* |
| Macrophage/Leukocytes | 91 | 41 | 132 | *mhc2a, coro1a, cxcr4b, mpeg1.1* |
| Leukocytes | 118 | 370 | 488 | *mhc2a, coro1a, cxcr4b* |
| Total | 3820 | 3657 | 7477 |  |

To determine clusters emerging upon injury, we assess the enriched cell composition for the injury. EC, Epi/cFB, MC, and throm were clearly distinguished by the injury status, identifying injury-induced subgroups (**Fig. 1B and Table 1**). We next focused on clusters enriched with *lepb* expression. *lepb* is detectable in the cells of injured but nearly undetectable in the uninjured hearts, indicating a regeneration-specific feature of *lepb* (**Fig. 1D and S1D**). The major clusters expressing *lepb* are the activated/injured EC. Interestingly, we found that *lepb* expression is high in some, but not all, activated EC clusters, revealing the heterogeneity of ECs in injured hearts. Although *lepb* expression in ECs was previously reported ^17, 21, 42^, our analysis identified additional *lepb* expressing cell type that one of the activated Epi/cFB clusters have the notable number of *lepb* expressing cells. This *lepb*^+^ Epi/cFB cluster displayed higher expression of *tcf21*, a well-defined Epi and epicardial-derived cell (EPDC) marker^43^, compared to another activated Epi/cFB cluster. In addition, some of the less well-defined clusters, including ECs containing uninjured cells and leukocytes, are combined. Collectively, our scRNA-seq analysis identified 15 distinct clusters representing various cardiac cell types and the injury-induced status (**Fig. 1A**).

### Identification of *lepb^+^* EC subgroup that directs injury-induced expression of secreted factors

The *lepb* expression level can separate the activated ECs into subgroups (**Fig. 2A**), prompting us to dissect the molecular basis of distinctly activated ECs. To end this, we analyzed differentially expressed genes between *lepb*^+^ and *lepb*^-^ activated ECs (**Supplementary Data S1**). Our analysis identified 80 genes with significant changes in expression levels (p-value <0.05 and fold change > 2), including 47 and 33 genes with increased and decreased expression, respectively. Gene Ontology (GO) analysis of these downregulated genes (representing *lepb^-^* activated ECs) indicated enrichment in regulation of transcription by RNA polymerase II and cell differentiation, while upregulated genes (representing *lepb^+^* activated ECs) were enriched for responses to stress, response to biotic stimulus, response to stimulus, and response to chemical (**Fig. S2A**). Consistent with GO analysis, gene set enrichment analysis (GSEA) demonstrated a significant increase of components for stress response (**Fig. S2B**). A major category of highly expressed genes in *lepb^-^* activated ECs is transcription factors (TFs), including the injury-responsive AP-1 complex (*fos* and *jun* family TFs)^44^ and endothelial/endocardial TFs (*klf2a* and *klf6a*)^45, 46^. However, further analysis to visualize cells expressing these factors demonstrated that a significant number of *lepb^+^* activated ECs expresses these TFs (**Fig. S2C**). A possible explanation is that these TFs are commonly expressed in both clusters, but the limited sequencing capacity causes biased reading. Another possible explanation is that the highly induced genes likely dampen the relative expression levels of these TFs in *lepb^+^* activated EC cluster.

**Figure 2.**
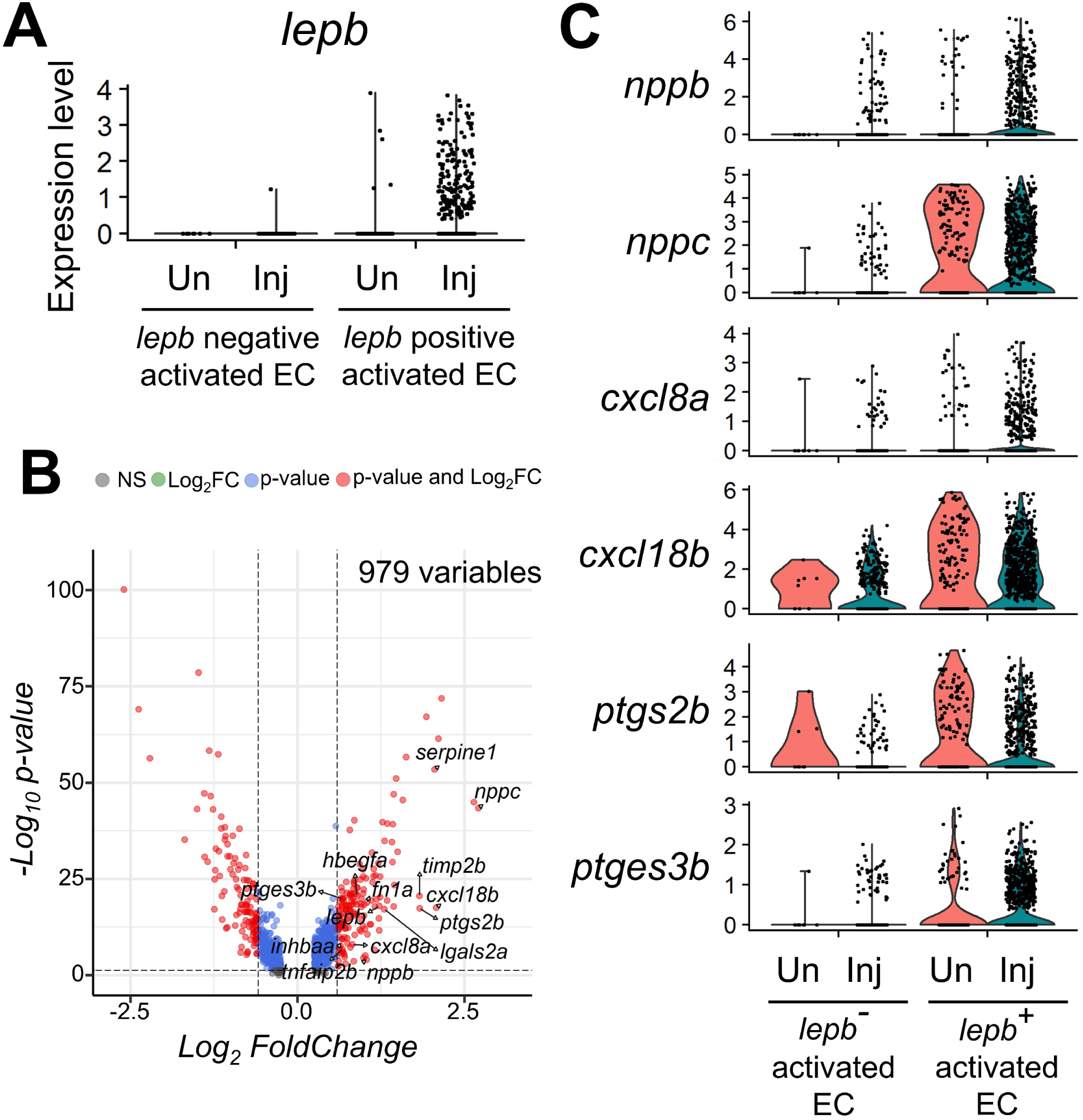
*lepb* marks a subset of EC population producing regenerative factors. (A) Robust induction of *lepb* in a subset of ECs activated by injury. (B) Differentially expressed genes between *lepb^+^* and *lepb^-^* activated ECs shown as a volcano plot. (C) Injury-dependent expression of regenerative factors generated by *lepb^+^* activated ECs. Inj, injured. Un, uninjured.

A sizable portion (18 of 47) of enriched genes (p-value <0.05 and fold change > 2) in *lepb^+^* activated ECs are secreted factors or genes related to synthesize secreted factors. *lepb^+^* activated ECs are also characterized by enrichment of regenerative factors or injury-inducible cytokines/chemokines, including *serpine1*^8^, *inhbaa*^47^, *fn1a*^26^, *hbegfa*^48^, *lgals2a*^49^*, timp2b*^50^*, tnfaip2b, cxcl8a*^51^, and *cxcl18b*^52^ (**Fig. 2B**, **2C and S2C**). Notably, two natriuretic peptides, *nppb* and *nppc*, are highly enriched in *lepb^+^* activated ECs; in fact, *nppc* was the most highly enriched gene in *lepb^+^* activated ECs. *nppb* and *nppc* are robustly induced in diseased and injured hearts^53–55^, providing additional evidence for *lepb^+^* ECs being an injury-activated cell population. Potent neutrophil recruitment chemokines, including *cxcl8a* and *cxcl18b*, were also highly enriched ^52, 56^, indicating that *lepb^+^* activated ECs produce chemokines to direct migration of immune cells to the wound site. We also noticed that two key prostaglandin E2 (PGE2) synthesis enzymes, including *prostaglandin-endoperoxide synthase 2b (ptgs2b,* also known as *cox2b)* and *prostaglandin E synthase 3b* (*ptges3b),* are enriched in *lepb^+^* activated ECs. PGE2 is an acute inflammatory signaling molecules that promote heart regeneration^57, 58^, implicating *lepb^+^* activated ECs as a source producing pro-regenerative factors. Therefore, our analysis highlights *lepb^+^* activated ECs as a signaling center that senses cardiac injury signals, produces signaling molecules to interact with other cardiac cells, such as immune cells, and secretes pro-regenerative factors to promote heart regeneration.

### Heterogenous Epi/EPDC lineages in regenerating hearts

Our scRNA-seq analysis defined two Epi/cFB clusters displaying signatures of cardiac fibroblasts, such as *col1a1a, col1a1b, col5a1, fn1a,* and *tagln*. As the number of cells derived from the uninjured hearts was considerably low (10% and 2%, **Table1**), these two clusters appear to emerge upon injury. In zebrafish, cFBs arise from *tcf21^+^* Epi upon injury^25, 59^, indicating that these two cFB clusters are derived from Epi. We analyzed major TFs governing central epicardial events, such as Epi formation, epicardial epithelial-to-mesenchymal transition (EMT), and EPDC lineage specification^60^ and found evident differences in the number of cells expressing *tcf21, tbx18a, snai2*, *twist1a*, *twist1b*, and *hand2.* Cells expressing these Epi-related TFs are high in the *tcf21^high^* Epi/cFB cluster (**Fig. 3B**). Notably, *tcf21^high^* Epi/cFB cluster also contains more *lepb* expressing cells than the *tcf21^low^* cluster (**Fig. 3B**). We next compared *tcf21^high^* and *tcf21^low^* clusters and identified 131 genes with significant changes in expression levels (p-value <0.05 and fold change > 2), including 63 and 68 genes with increased and decreased expression, respectively (**Fig. 3C and Supplementary Data S2**). GO analysis of the significantly downregulated genes (enriched in the *tcf21^low^* activated Epi/cFBs) indicated enrichment in regulation of localization, ion transmembrane transport, and cell morphogenesis involved in differentiation, whereas upregulated genes (enriched in the *tcf21^high^* activated Epi/cFBs) were associated with cell adhesion, defense response, wound healing, ECM, response to hormone (**Fig. S3A**). Consistent with GO analysis, GSE analysis indicated a significant increasement of components for cell adhesion, ECM organization and wound healing in *tcf21^high^* activated Epi/cFBs (**Fig. S3B**). Thus, our data suggested that two molecularly distinct subpopulations are present in the Epi/EPDC lineage upon injury.

**Figure 3.**
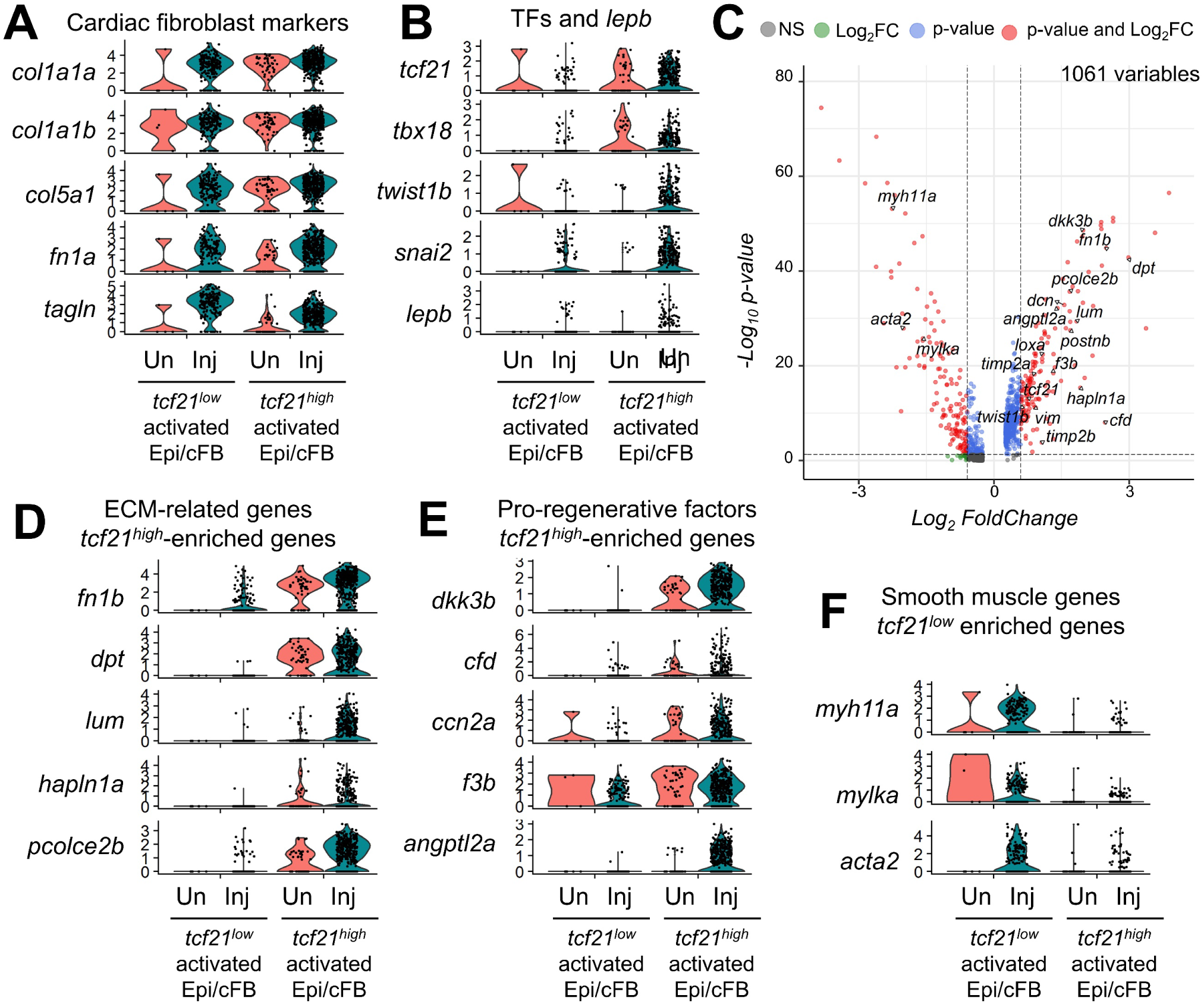
Heterogenous Epi/cFB in zebrafish injured hearts. (A) Injury-inducible expression of cardiac fibroblast marker genes in *tcf21^low^* and *tcf21^high^* activated Epi/cFB. (B) Enrichment of cells expressing key epicardial transcription factors (TFs) and *lepb* in *tcf21^high^* activated Epi/cFB. (C) Differentially expressed genes between *tcf21^low^* and *tcf21^high^* activated Epi/cFB shown as a volcano plot. (D, E) Highly expressed ECM-related genes (D) and pro-regenerative factors (E) in *tcf21^high^* activated Epi/cFB. (F) Highly expressed smooth muscle genes in *tcf21^low^* activated Epi/cFB.

The majority of enriched genes (p-value <0.05 and fold change > 2) in *tcf21^high^* activated Epi/cFBs consists of ECM-related genes, including *fn1b*^26^*, dpt*^61^*, dcn*^62^*, hapln1a*^63^*, postnb*^10^*, pcolce2b*^64^*, lum*^65^*, loxa*^66^*, mxra8a, vim*^67^*, timp2a,* and *timp2b*^68^ (**Fig. 3C**). Higher expression of ECM genes indicates that *tcf21^high^* activated Epi/cFBs are the major cell-type producing ECM and ECM-modifying enzymes upon injury, implying that this cluster potentially remodels extracellular environments in a manner favorable for heart regeneration. *tcf21^high^* activated Epi/cFBs are also characterized by several signaling factors, including *dkk3b* (wnt antagonists)^7, 69^, *cfd* (adipsin)^70^, and *ccn2a*^71^, and angiogenic factors, including *f3b* (coagulation factor III, *cd142b*)^72^ and *angptl2a*^73^. These factors are considered to be pro-regenerative factors in injured tissues, implicating the paracrine signaling roles of *tcf21^high^* activated Epi/cFBs. In contrast, *tcf21^low^* activated Epi/cFBs are enriched with smooth muscle markers, such as *myh11a, mylka* and *acta2,* indicating that these cells are likely vascular smooth muscle cells derived from epicardium. Thus, our analysis indicates that Epi/cFBs exhibit heterogeneity upon injury with a unique molecular feature of key TF and *lepb* expressing subpopulation.

Sun and colleagues recently published a report to analyze *tcf21^+^* sorted epicardial cells at the single cell level, demonstrating that Epi/EPDCs are the heterogenous populations in regenerating zebrafish hearts^15^. As their scRNA-seq analysis used substantial numbers of epicardial cells (around 4,000 cells for each uninjured and regenerating hearts), we utilized this high-quality scRNA-seq data to further dissect *lepb* enriched *tcf21^+^* Epi/EPDCs. *lepb^+^ tcf21^+^* clusters are expected to be unique based on the fact of low numbers of *lepb^+^ tcf21^+^* cells, and thus we increased UMAP resolution to separate Epi/EPDC clusters explicitly and identified 16 clusters (**Fig. 4A and Table S1**). Similar to Sun and colleagues report, we found cell cycle gene-enriched cluster (Cluster 13 characterized with *mki67, fen1, mcm2, PCNA,* and *rpa2*), defense-responsive cluster (Cluster 6 characterized with *mxb, rsad2* and *saa*), *crabp1a^+^* and *Frzb^+^* cluster (Clusters 8 and 14) and *cxcl12b^+^* cluster (Cluster 12) (**Fig. S4**). Our analysis subdivided the largest cell-contained and immune responsive clusters in the original report into multiple clusters. Interestingly, *lepb* expression is higher in regenerating samples of the Cluster 10 (**Fig. 4B and C**). This *lepb^+^* Epi/EPDC cluster is characterized with chemokine and inflammation-related genes, such as *cxcl8b.1*^74^, *c3a.3*^75^, and *steap4*^76^ (**Fig. 4C**), postulating their roles in the immune response. Overall, our approach demonstrates that scRNA-seq analysis combined with regeneration-specific gene profiles can subset the heterogenous clusters to identify unique subpopulations.

**Figure 4.**
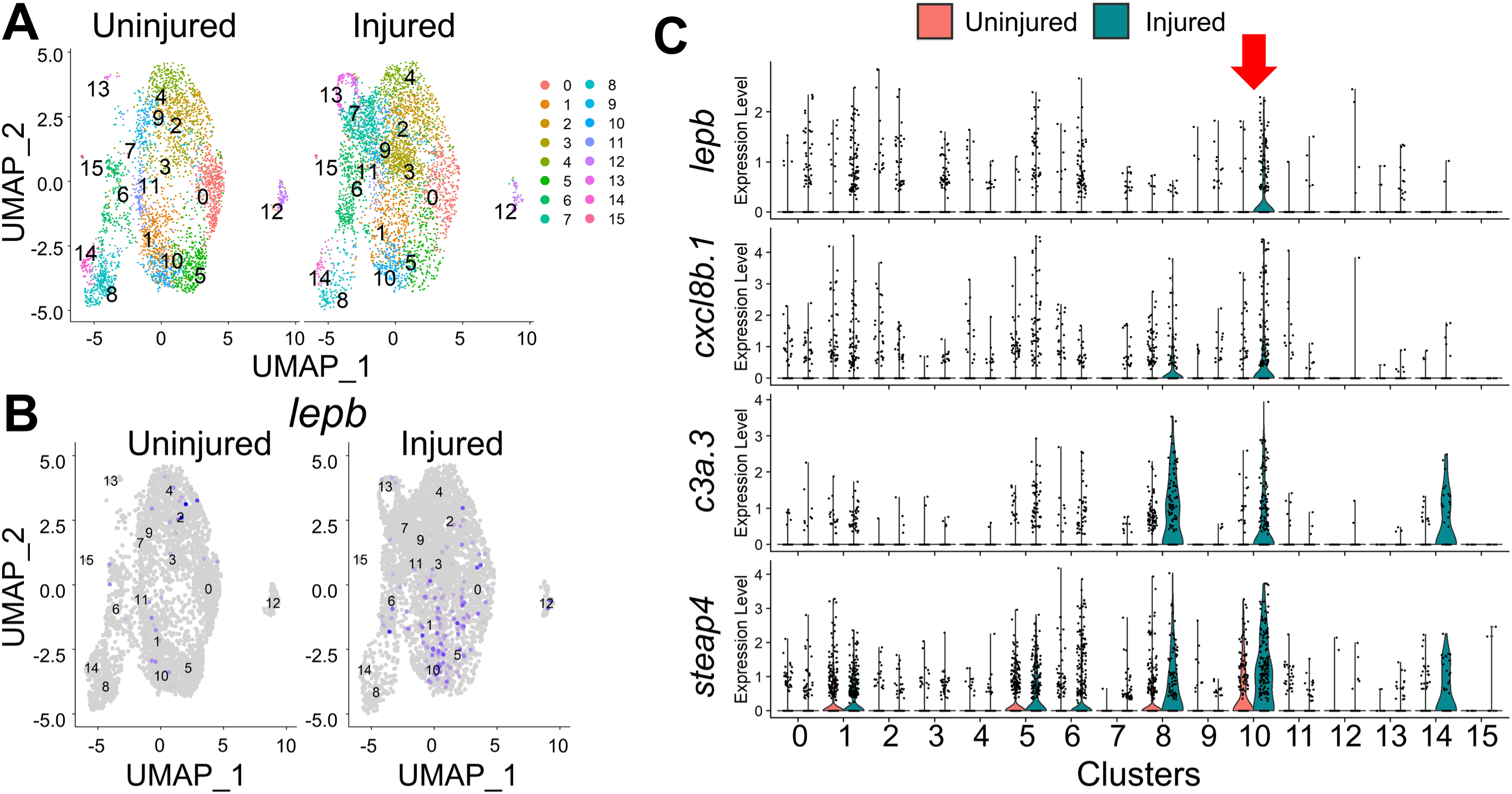
*lepb* marks a specific subpopulation of *tcf21^+^* Epi/EPDCs. (A) Clustering assignments of *tcf21^+^* Epi/EPDCs collected from uninjured and injured hearts. (B) Injury-dependent expression of *lepb* depicted by UMAP plot. (C) Enrichment of cells expressing the Cluster 10 specific genes (TFs), including *lepb, cxcl8b.1, c3a.3* and *steap4*.

### Epi/cFB-specific epigenomic profiles indicate *LEN* is active in a subset of Epi/cFBs upon injury

*LEN* was identified as an enhancer directing regeneration-specific expression in hearts, which is regulated by a ∼300 bp sequence at the proximal end of *LEN* (cardiac *LEN* or *cLEN*)^17^. Our previous work demonstrated that *cLEN* can direct injury-induced expression in a subset of EC ^21^, which is likely the *lepb^+^* EC cluster defined by our scRNA-seq analysis as *lepb* is the target gene of *LEN*. scRNA-seq analysis also demonstrated *lepb* expressing cells in *tcf21^high^* Epi/cFBs and the Cluster 10 of *tcf21^+^* Epi/EPDCs, but *LEN* activity in Epi/EPDCs has not been assessed. We first examined *lepb* expression levels with RNA-seq profiles of Epi/EPDCs generated by *tcf21:EGFP^+^* cells sorted from 0, 3 and 7 days post amputation (dpa) hearts ^20^. Compared to the 0 dpa sample, *lepb* RNA levels were sharply elevated at 3 and 7 dpa with a 2.69 and 3.86 fold change (**Fig. 5A**), respectively, indicating injury-inducible *lepb* expression. To determine whether *LEN* is accessible in Epi/EPDCs upon injury, we used Assay for Transposase-Accessible Chromatin using sequencing (ATAC-seq) of *tcf21:EGFP^+^* cells sorted from 0, 3 and 7 dpa hearts. ATAC-seq profiles of *tcf21^+^* cells showed *LEN* accessibility is increased by 1.62 fold in 3 dpa samples (FDR = 0.001008207), compared to 0 dpa (**Fig. 5A**), indicating that *LEN* is activated in Epi/EPDC cells upon injury.

**Figure 5.**
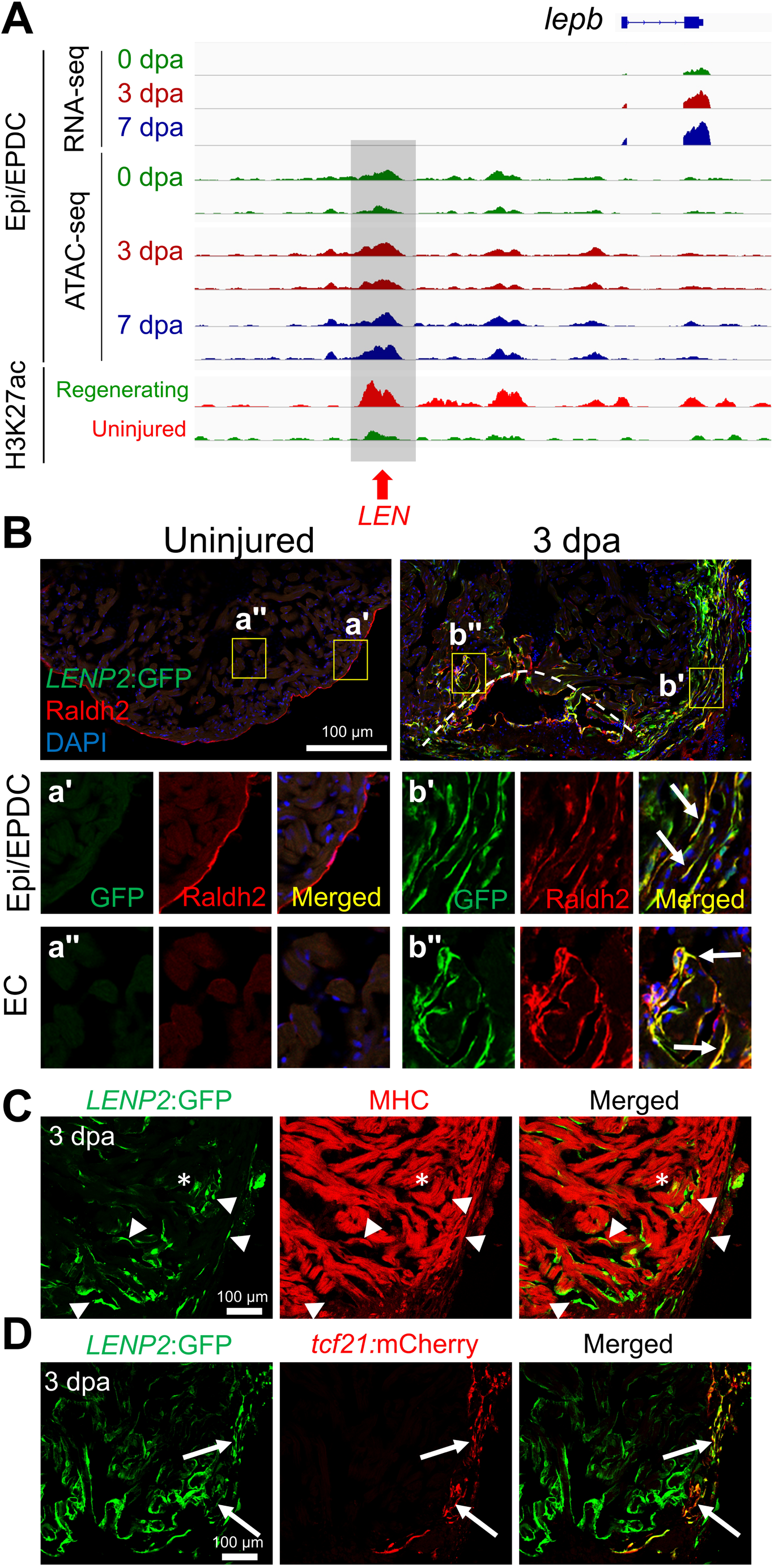
*LEN* directs injury-induced Epi/EPDC expression. (A) Genome browser tracks of the genomic region near *lepb* showing the transcripts and chromatin accessibility profiles in the Epi/EPDC. The whole-ventricle H3K27Ac profile of the uninjured and regenerating heart is shown at the bottom. Gray box and red arrow indicate *LEN*. (B-D) Immunostained section images of transgenic fish carrying *LENP2:EGFP*. (B) Raldh2 antibody is used to label EC and Epi/EPDC. Uninjured heart shows one single Raldh2^+^ cell layer outlining the cardiac chamber. Raldh2 signal emerges in the ECs at the wound area and EPDCs in the cortical layers upon injury, which are co-labelled with *LENP2:EGFP* (Arrows). The boxed areas are enlarged at the bottom panels. (C) Myosin heavy chain (MHC) antibody is used to label CMs. (D) *tcf21:mCherry* is used for Epi/EPDC expression. While *LENP2:EGFP* rarely colocalizes with MHC^+^ CMs (Arrowheads), a subset of *LENP2:EGFP* co-localizes with *tcf21:mCherry* (Arrows). Note that asterix indicates CM expression as a basal expression of the P2 minimal promoter. At least five hearts for uninjured and injured samples were examined and all animals displayed a similar expression pattern.

We next asked whether *LEN* can direct injury-dependent expression in Epi/ EPDC *in vivo*. *LENP2:EGFP* transgenic fish carry enhancer reporter constructs, in which *LEN* is coupled with the 2 kb upstream minimal promoter of *lepb* (*P2*). To assess injury-dependent activity, we amputated the apex of ventricles, collected hearts at 3 dpa, and immunostained cardiac section samples with a Raldh2 antibody. While Raldh2 was detected only in the outermost Epi in the uninjured hearts, cardiac injury expanded Raldh2 expression to the EC, Epi and EPDCs^5, 77^. *LENP2:EGFP* has no EGFP expression in uninjured adult hearts but are robustly induced EGFP at 3 dpa (**Fig. 5B**). As previously reported^17, 21^, we observed strong EGFP signals in Raldh2^+^ EC cells near the amputation site and a subset of EC cells inside of the ventricle at 3 dpa (**Fig. 5B**). In uninjured hearts, *tcf21^+^* Epi cells are restricted to the outermost ventricular layer, but a subset of *tcf21^+^* Epi emerges in the cortical myocardial layers to generate cFBs and other EPDC lineages^25^. We identified that *LENP2:EGFP* are also highly detectable in Raldh2^+^ and *tcf21:mCherry^+^* Epi/EPDCs in the cortical muscle of ventricles, confirming *LEN* activity in Epi/EPDC (**Fig. 5B-D**). Thus, our results demonstrate that *LEN* drives injury-dependent expression in EC and Epi/EPDCs, two major non-cardiomyocyte cells.

### *LEN* deletion abolishes injury-dependent *lepb* induction in the hearts, but not other surrounding genes

Recent advent of genome editing allows to delete enhancers to assess their functional significance. To determine whether *LEN* is required for injury-induced *lepb* expression in hearts, we established *LEN* deletion mutants (*LEN^Δ^*), in which 3.7 kb surrounding *LEN* is removed by CRISPR/Cas9 (**Fig. S5**)^78^. *LEN* deletion animals are viable, and we did not detect any noticeable overt phenotype, such as obesity, under the standard husbandry condition. To examine whether *LEN* deletion causes defects in heart regeneration, we quantified CM proliferation at 7 dpa and fibrotic scar resolution at 30 dpa after partial ventricular resection. *LEN^Δ/Δ^* mutants exhibited normal injury-induced CM proliferation and were able to resolve fibrotic scar similar to that of controls (**Fig. 6A-C**). These results are in agreement with our previously discovery that *lepb* mutants regenerate heart normally ^17^.

**Figure 6.**
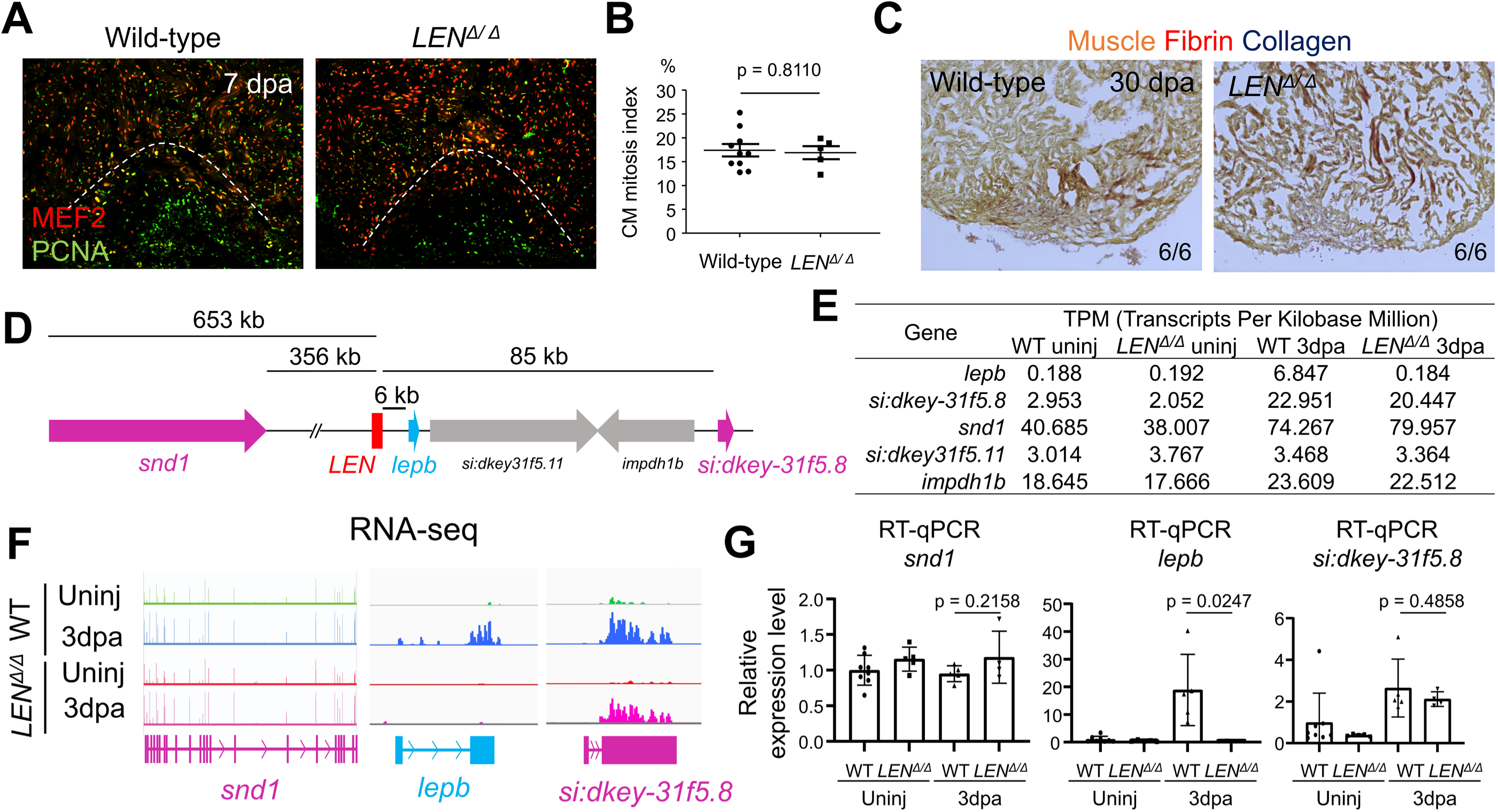
*LEN* is essential for injury-dependent *lepb* induction during heart regeneration. (A) Representative images of PCNA/Mef2 staining quantified in (B). (B) Quantification of adult ventricular CM proliferation indices for 7 dpa wild-type sibling (control) and *LEN^Δ/Δ^* hearts. n = 10 and 5 for control and *LEN^Δ/Δ^* hearts. 3 sections per heart were used. (C) Representative images of AFOG staining for 30 dpa control and *LEN^Δ/Δ^* hearts. n = 6 for control and *LEN^Δ/Δ^* hearts. 3 sections per heart were used. (D) Genomic region surrounding *LEN*. *snd1, lepb, si:dkey-31f5.8* and *rap1b* are induced during heart regeneration. (E) Transcripts Per Kilobase Million (TPM) for *lepb* and its surrounding genes. (F) RNA-seq of uninjured and 3 dpa wild-type (WT) and *LEN^Δ/Δ^* hearts. Tracks indicate an absence of *lepb* expression at 3 dpa in *LEN^Δ/Δ^* fish, whereas *snd1* and *si:dkey-31f5.8* transcript levels are increased similarly in both wild type and *LEN^Δ/Δ^* upon heart injury. (F) RT-qPCR analysis of *snd1, lepb,* and *si:dkey-31f5.8* transcript levels in WT and *LEN^Δ/Δ^* hearts. *lepb* is undetectable in 3 dpa injured *LEN^Δ/Δ^* hearts. unpaired two-tailed t-tests were performed to indicate significance. (n=8, 5, 5, and 4 for WT uninjured, *LEN^Δ/Δ^* uninjured, WT 3dpa, and *LEN^Δ/Δ^* 3dpa, respectively). Data are mean ± s.d.

To examine the requirement of *LEN* in control of endogenous *lepb* expression upon injury, we performed RNA-seq analysis with uninjured and 3 dpa hearts of control and *LEN^Δ/Δ^* mutants. *lepb* transcript level is sharply upregulated with a 36-fold increase at 3 dpa control hearts compared to uninjured control hearts (**Fig. 6D-F**). Notably, transcriptome analyses revealed no detectable elevation of *lepb* at 3 dpa of *LEN^Δ/Δ^* hearts (**Fig. 6E, F**). These results suggest that injury-induced *lepb* expression is governed by *LEN* and there is no alternative enhancer element to redundantly regulate *lepb* expression at this locus. To determine whether *LEN* can control injury-dependent expression of multiple genes, we surveyed expression of neighboring genes, including *si:dkey-31f5.8* (85kb downstream), and *snd1* (359kb upstream). Injury-dependent gene expression of these genes was hardly affected by *LEN* deletion in 3 dpa hearts (**Fig. 6D-F**). RNA-seq analysis demonstrated that *si:dkey-31f5.8* and *snd1* are induced upon injury in control hearts with 7.77 and 1.83 fold change, respectively. In *LEN^Δ/Δ^* hearts, these genes are also induced upon injury with 9.96 and 2.10 fold change, respectively (**Fig. 6E, F**). *si:dkey31f5.11* and *impdh1b,* other two genes in close proximity to *lepb*, are not differentially expressed upon injury, indicating they are not cardiac injury-responsible genes (**Fig. 6E**). Next, we validated our RNA-seq results using quantitative reverse transcription PCR (RT-qPCR) analysis. Our RT-qPCR analysis also demonstrated that *si:dkey-31f5.8* was upregulated at 3 dpa of WT and *LEN^Δ/Δ^* hearts and that robust *lepb* induction was abrogated in *LEN^Δ/Δ^* , but not WT, hearts (**Fig. 6G**). *snd1* was not upregulated in both WT and *LEN^Δ/Δ^* in our RT-qPCR analysis (**Fig. 6G**). Collectively, our results demonstrate that *LEN* specifically governs injury-induced *lepb* expression, but not other neighboring genes.

## DISCUSSION

The heart is a complex tissue comprising of multiple cell types, of which intercellular interaction is crucial for heart repair. Here, we analyzed scRNA-seq data of uninjured and injured hearts and utilized *lepb*, a regeneration-specific gene, to enhance analysis power to identify unique cell subpopulations. Interestingly, *lepb* specifies EC and Epi/EPDC linages into distinct populations in the injured hearts. The prominent feature of *lepb^+^* ECs is the over- representation of secreted factors, which are known to act as regenerative factors. In zebrafish hearts, ECs are the most rapidly responding cell types to injury cues by changing their morphology and activating expression of secreted factors within hours of injury. As ECs are in direct contact with CMs, ECs are considered to be the most effective cell types to interact with CMs ^5, 79^. *LEN* activity is restricted to the wound area^21^, suggesting that *lepb^+^* ECs emerge at the wound area upon injury and serve as a paracrine signaling center to trigger CM proliferation. *lepb^+^* ECs are enriched with *cxcl8a* and *cxcl18b* (immune cell attractant chemokines^52, 56^), and *lepb*-enriched Epi/EPDCs are characterized with high expression of *cxcl8b.1*, *c3a.3,* and *steap4* (immune-related genes^74–76^), highlighting their roles for immunomodulation at the wound area.

*Leptin*, encoded by the *obese* (*ob*) gene, is a well-characterized adipocytokine controlling feeding and energy balance regulation^80^. In addition to these obesity-related effects, multiple studies demonstrated regeneration-associated roles of *Leptin*. In mammals, *Leptin* levels are elevated in cardiovascular disease like myocardial infarction (MI)^81, 82^ and in skin upon injury^83^, revealing *Leptin* as an injury-inducible factor across vertebrates. *Leptin* also exerts pro-regenerative functions in multiple tissues, including skin and hearts^82–86^. For instance, *Leptin* mutant mice (*ob*) showed higher mortality after cardiac injury, whereas administration of Leptin in *ob* mice yielded improved cardiac function and survival rate^82, 84, 85^. Zebrafish have two *leptin* homologs: *lepa* and *lepb*. Although roles for zebrafish *leptin* signaling in the regulation of the feeding and obesity are unclear due to contradicting observation of the obese phenotype^87, 88^, their involvement in tissue regeneration has been suggested. In eyes, *lepa* and *lepb* are robustly induced upon injury and administration of Leptin can stimulate eye regeneration through the Jak/Stat pathway^89^. The same study also revealed that *il6* family cytokines, including *il11* and *cntf*, were able to stimulate eye regeneration via the Jak/Stat pathway. While *LEN* deletion and *lepb* mutant^17^ fish can regenerate their hearts, *lepb* regenerative roles may be compensated by *il11* signaling as both *il11* and *leptin* signaling share downstream effectors, such as Jak/Stat. Our transcriptomic analysis indicates that *lif/m17*, *il11* family gene, is upregulated upon injury in both control and *LEN^Δ/Δ^* hearts (**Fig. S6**). Recent study reported that *il11 receptor a* (*il11ra*) mutant fish displayed impaired heart regeneration by failed scar resolution and decreased CM proliferation at the later phase of hear regeneration^90^. However, this work indicated that CM proliferation at 7 dpa is likely normal, suggesting that *il11* signaling on CM proliferation at the early phase of heart regeneration may be compensated by Leptin signaling.

TREEs are crucial for triggering injury-dependent expression in a tissue-specific manner and directing gene expression stably during regeneration. Much research in recent years have been performed to identify TREE or regeneration enhancer candidates using genome-wide analysis. *in vivo* activity of several candidates, including *LEN*, have been confirmed via transgenic assays^4, 19, 20, 78, 91^. In addition to the typical transgenic assay, we validated the *LEN* activity directing regeneration-dependent gene expression using the enhancer deletion line. Importantly, we demonstrated that *LEN* deletion completely abrogated injury-inducible expression of *lepb* in hearts without affecting other nearby genes. This implies that one class of TREEs selectively regulates expression of a single gene within a short range. Exploring 3D chromatin conformational change of this short-ranged TREE and nearby regions upon injury will be interesting future work to understand how 3D genome architecture change affects regeneration-dependent transcription.

Heart regeneration is a highly complicated process governed by diverse cell populations with various transcriptional programs. Our integrative analyses of multiple sequencing data and genetic animal models characterize regeneration-emerging cell types and regulatory elements, leading to discovery of novel regeneration features, such as cells, factors, and *cis*-regulatory elements.

### Zebrafish maintenance and procedures

Wild-type or transgenic male and female zebrafish of the outbred Ekkwill (EK) strain ranging up to 18 months of age were used for all zebrafish experiments. The water temperature was maintained at 26°C for animals unless otherwise indicated. Partial ventricular resection surgery was performed as described previously ^92^, in which ∼20% of the cardiac ventricle was removed at the apex. For expression patterns to determine enhancer activity, at least four hearts were examined per experiment. To define *LEN* and *cLEN* activity, *Tg(LENP2:EGFP)^pd1^*^30^ and *Tg(tcf21:mCherry-NTR)^pd108^* were used. *LENΔ/Δ^pd281^* was created using CRISPR/Cas9 as described in ^78^ using acgATTTAGGTGACACTATAGAatgtatccgtataccata GTTTTAGAGCTAGAAAtagc and acgATTTAGGTGACACTATAGAgaacccaattaggattta GTTTTAGAGCTAGAAAtagc oligos. Work with zebrafish species was performed in accordance with University of Wisconsin-Madison guidelines.

### Histology and imaging

Hearts were fixed with 4% paraformaldehyde overnight at 4°C or for 1 h at room temperature. Cryosectioning and immunohistochemistry were performed as described previously ^21^. Hearts were cryosectioned at 10 µm thickness. Heart sections were equally distributed onto four or five serial slides such that each slide contained sections representing all areas of the ventricle. A solution comprising 5% goat serum, 1% bovine serum albumin, 1% dimethyl sulfoxide and 0.1% Tween-20 was used for blocking and antibody staining. The primary and secondary antibodies used in this study were as follows: anti-myosin heavy chain (mouse; F59; Developmental Studies Hybridoma Bank; 1:50), anti-EGFP (rabbit; A11122; Life Technologies; 1:200), anti-EGFP (chicken; GFP-1020; Aves Labs; 1:2000), anti-Ds-Red (rabbit; 632496; Clontech; 1:500), anti-Raldh2 (rabbit; GTX124302; Genetex; 1:200), anti-MHC (mouse; F59; Developmental Studies Hybridoma Bank), Alexa Fluor 488 (mouse, rabbit and chicken; A11029, A11034 and A11039; Life Technologies; 1:500) and Alexa Fluor 594 (mouse and rabbit; A11032 and A11037; Life Technologies; 1:500). anti-MEF2 (rabbit, sc- 313, Santa Cruz Biotechnology), anti-PCNA (mouse, P8825, Sigma), Alexa Fluor 488 (mouse and rabbit; Life Technologies), Alexa Fluor 594 (mouse and rabbit; Life Technologies). Cardiac section images were acquired using BZ-X810 fluorescence microscope (Keyence), LSM 700 confocal microscope (Zeiss), A1R-s confocal microscope (Nikon). Image stitching was automatically processed using BZ-X800 analyzer. Further image processing was carried out manually using Photoshop or FIJI/ImageJ software. For AFOG staining, cardiac cryosection slides were fixed with Bouin’s solution for 2 hours at 60°C and stained as described previously ^92^. Imaging was performed using Eclipse Ti-U inverted compound microscope (Nikon) and processed by Photoshop.

### RNA isolation and qPCR

RNA was isolated from uninjured and partly resected hearts using TriReagent (ThermoFisher). Complementary DNA (cDNA) was synthesized using a NEB ProtoScript II first strand cDNA synthesis kit (NEB, E6560). Quantitative PCR was performed using the qPCRBIO SyGreen Blue Mix Separate-ROX (Genesee Scientific, 17-507) and a Bio-Rad CFX Connect system. All samples were analyzed in at least biological quadruplicate with two technical repeats. The sequences of the primers used are listed in Table S1. Transcript levels were normalized to *actb2* levels in all experiments.

### RNA-seq and ATAC-seq analyses

For RNA-sequencing, total RNA was prepared from uninjured and 3 dpa resected hearts of wild-type siblings and *LEN^Δ/Δ^*. Generation of mRNA libraries and sequencing were performed at the Duke Center for Genomic and Computational Biology using Illumina HiSeq4000 with 50bp single read runs. Adapter sequences were trimmed by Cutadapt. Sequences were aligned to the zebrafish genome (genome assembly GRCz11, Ensembl gene annotation release 104) using HISAT2^93^. Gene counts were obtained by featureCounts and Transcripts Per Kilobase Million (TPM) was used to calculate fold-change. For epicardium, we used RNA-seq and ATAC-seq datasets of GSE89444, which were aligned to the GRCz11 using HISAT2. IGV genome browser was used to browser track images. Accession numbers for transcriptome data sets are GSE199697.

### scRNA-seq analysis

For scRNA-seq analysis of uninjured and injured hearts, we obtained original sequencing files from GSE138181 ^22^ and reanalyzed this profile using 10x Genomics cloud service (https://cloud.10xgenomics.com/cloud-analysis). The Danio_rerio.GRCz11 (release 104) version of the zebrafish reference genome and annotation files were downloaded from Ensemble database (ensembl.org). Raw counts of wild-type uninjured and injured hearts were used for scRNA-seq analysis with the Seurat package^94^. Low quality cells (nUMI ≤ 500, nGene ≤ 250, mitoRatio > 0.15, log_10_GenesPerUMI < 1.7) were filtered out. After careful inspection, the 40 principal components (PCs) of the PCA with resolution 1 were used for clustering. Differential expression (DE) analysis was performed using the FindMarkers function of the Seurat package. GO-term and GSEA analyses were done by the enrichGo and gseGO functions of clusterProfiler^95^. Volcano plot was generated by the EnhancedVolcano package ^96^.

For scRNA-seq analysis of Epi/EPDC cells, we obtained raw count files from GSE172511^15^ and reanalyzed it. Low quality cells (nUMI ≤ 500, nGene ≤ 250, mitoRatio > 0.15, log_10_GenesPerUMI < 1.0) were filtered out. Nonepicardial cells (*myl7*, *fli1a*, or *lcp1* positive cells) and *tcf21^-^* cells were filtered out. After careful inspection, the 40 principal components (PCs) of the PCA with resolution 1.4 and integration were used for clustering using the Seurat package^94^.

## DATA AND MATERIALS AVAILABILITY

Sequencing data have been deposited in GEO under accession code GSE199697. Reviewer token is kjstqswcbrgpncz.

## ACKNOWLEDGMENTS

We thank the UW-Madison SMPH BRMS staff for zebrafish care; Nutishia Lee for contributions to experiments; Shuyang Chen for comments on bioinformatic analysis; Kang lab members for comments on the manuscript.

## AUTHOR CONTRIBUTIONS

Biological experiments: KS, IJB, JK Computational analysis and data curation: IJB, JK Conceptualization: JK Writing original draft: JK Reviewing, Editing: KS, IJB, JC, JK Supervision: JK Funding: JC, JK

## DECLARATION OF INTERESTS

The authors declare no competing interests.

## FUNDING

AHA AHA16SDG30020001 (JK)

R01 National Institutes of Health grant R01 HL151522 (JK)

National Institutes of Health, University of Wisconsin Carbone Cancer Center Support grant P30 CA 014520 (JK)

National Heart Lung and Blood Institute Training grant T32 HL 007936 (IJB)

National Heart Lung and Blood Institute Training grant F31 HL 162492 (IJB)

American Heart Association Predoctoral Fellowship 827904 (IJB)

AHA Career Development Award, 18CDA34110108 (JC)

National Heart Lung and Blood Institute R01 HL155607 (JC)

## SUPPLEMENTARY MATERIAL

**Supplementary Figures S1-S6**

**Supplementary Table S1. Cell number of tcf21+ Epi/EPDC clusters in the uninjured and injured hearts identified by scRNA-seq analysis**

**Supplementary Data S1. Differentially expressed gene list in *lepb^+^* and *lepb^-^* ECs.**

**Supplementary Data S2. Differentially expressed gene list in *tcf21^high^* and *tcf21^low^* Epi/cFBs.**

---

**Figure S1.**
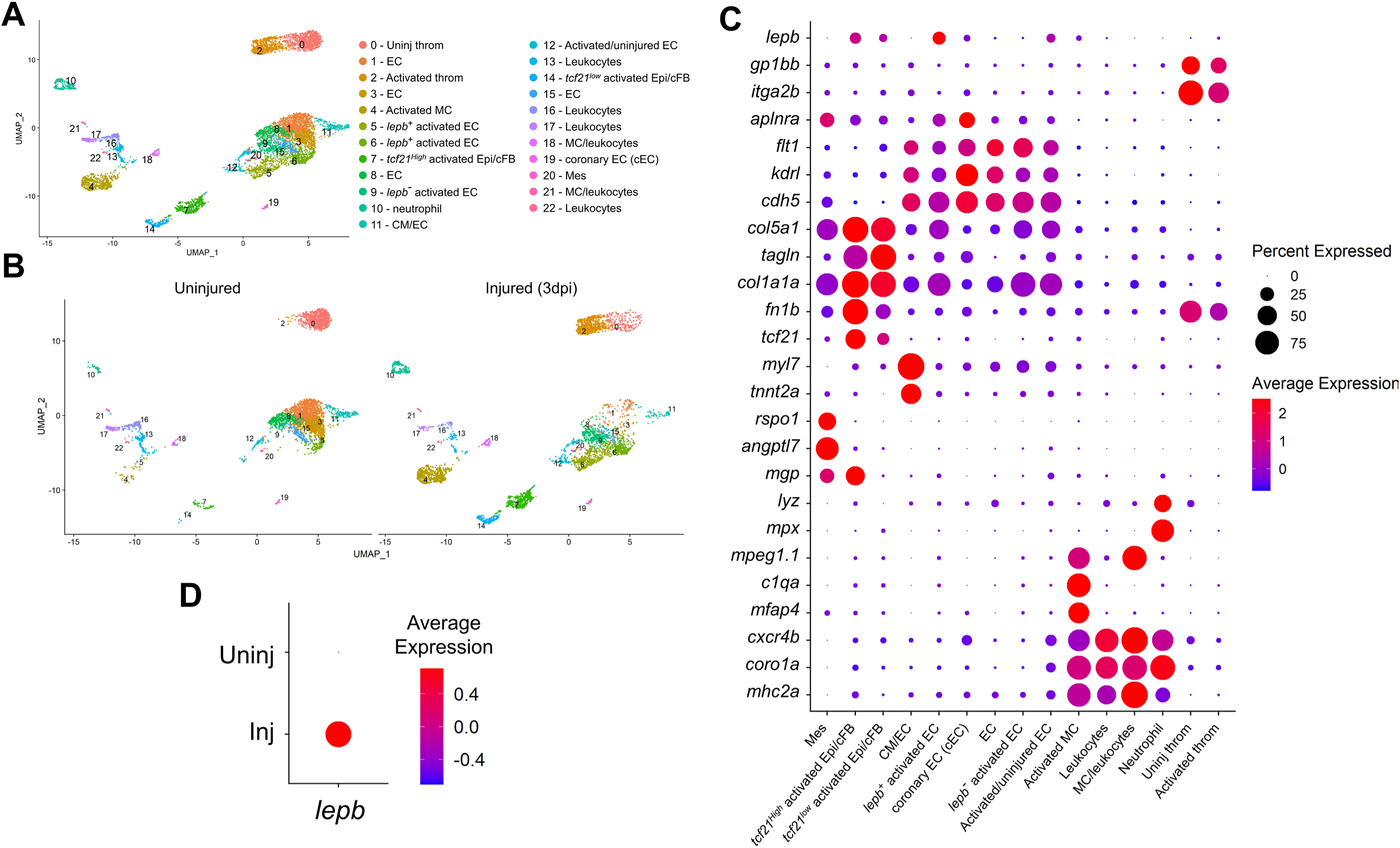
Heterogenous cell clusters of adult zebrafish. (A) Raw data of clustering assignments of *runx1P2:Citrine* or *kdrl:mCherry* expressing cells collected from uninjured and injured hearts. In Fig. 1A, the less well-defined clusters, including ECs, leukocytes, are combined. throm, thrombocyte. EC, endocardial/endothelial cells. Epi, epicardial cells. cFB, cardiac fibroblasts. Mes, mesenchyme-like cells. MC, macrophages. (B) Clustering assignments for uninjured and injured hearts. dpi, days post-injury. (C) Differential expression of the key marker genes to identify cell types shown as a dot plot. (D) Injury-dependent expression of *lepb* depicted by a dot plot.

**Figure S2.**
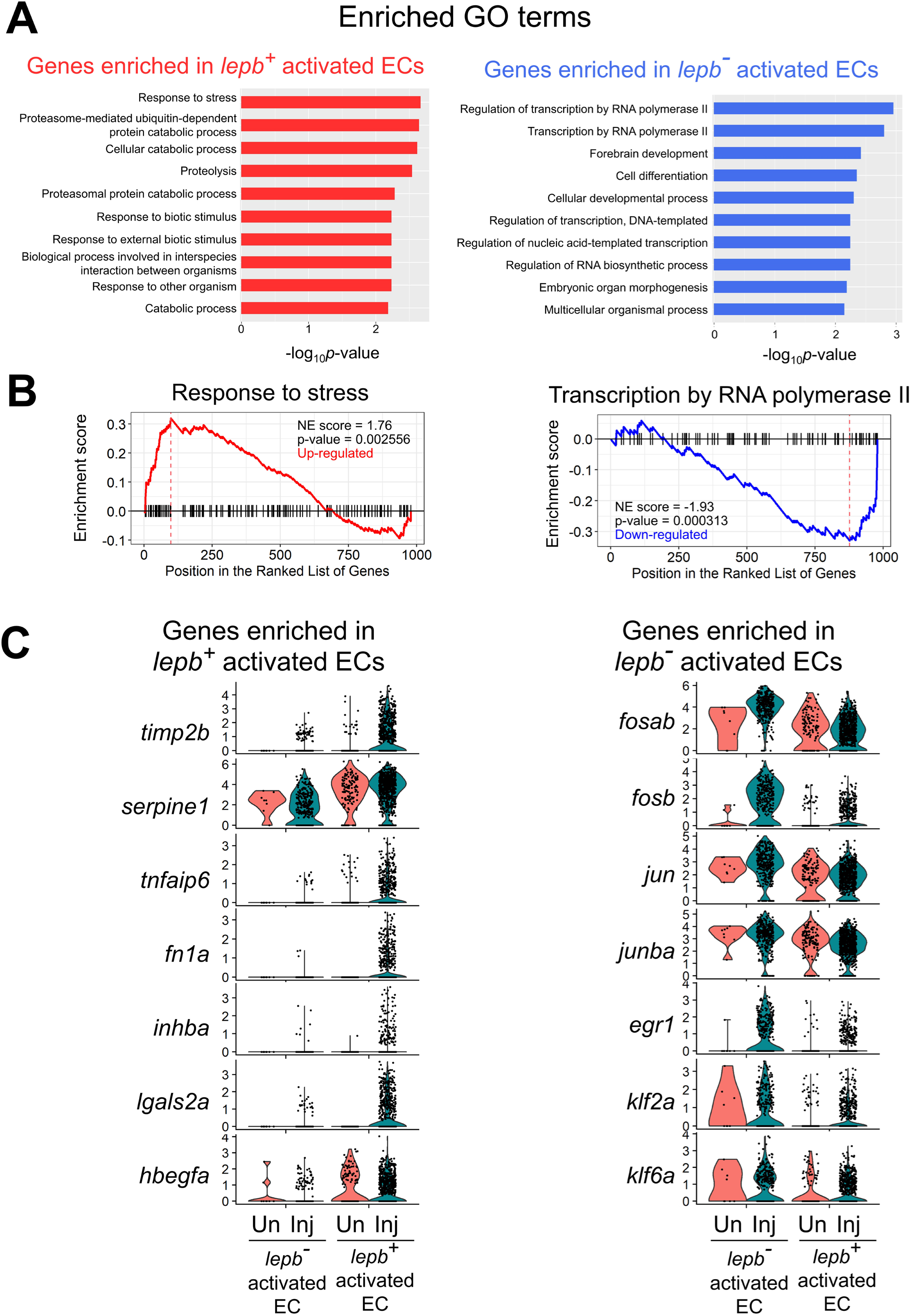
Differential gene expression analysis between *lepb^+^* and *lepb^-^* activated ECs. (A) Top GO terms for genes enriched in *lepb^+^* (Red) and *lepb^-^* (Blue) activated ECs. (B) GSE analysis plot of response to stress and transcription by RNA polymerase II. NE score, normalized enrichment score. (C) Genes enriched in *lepb^+^* and *lepb^-^* activated ECs shown as a violin plot. Inj, injured. Un, uninjured.

**Figure S3.**
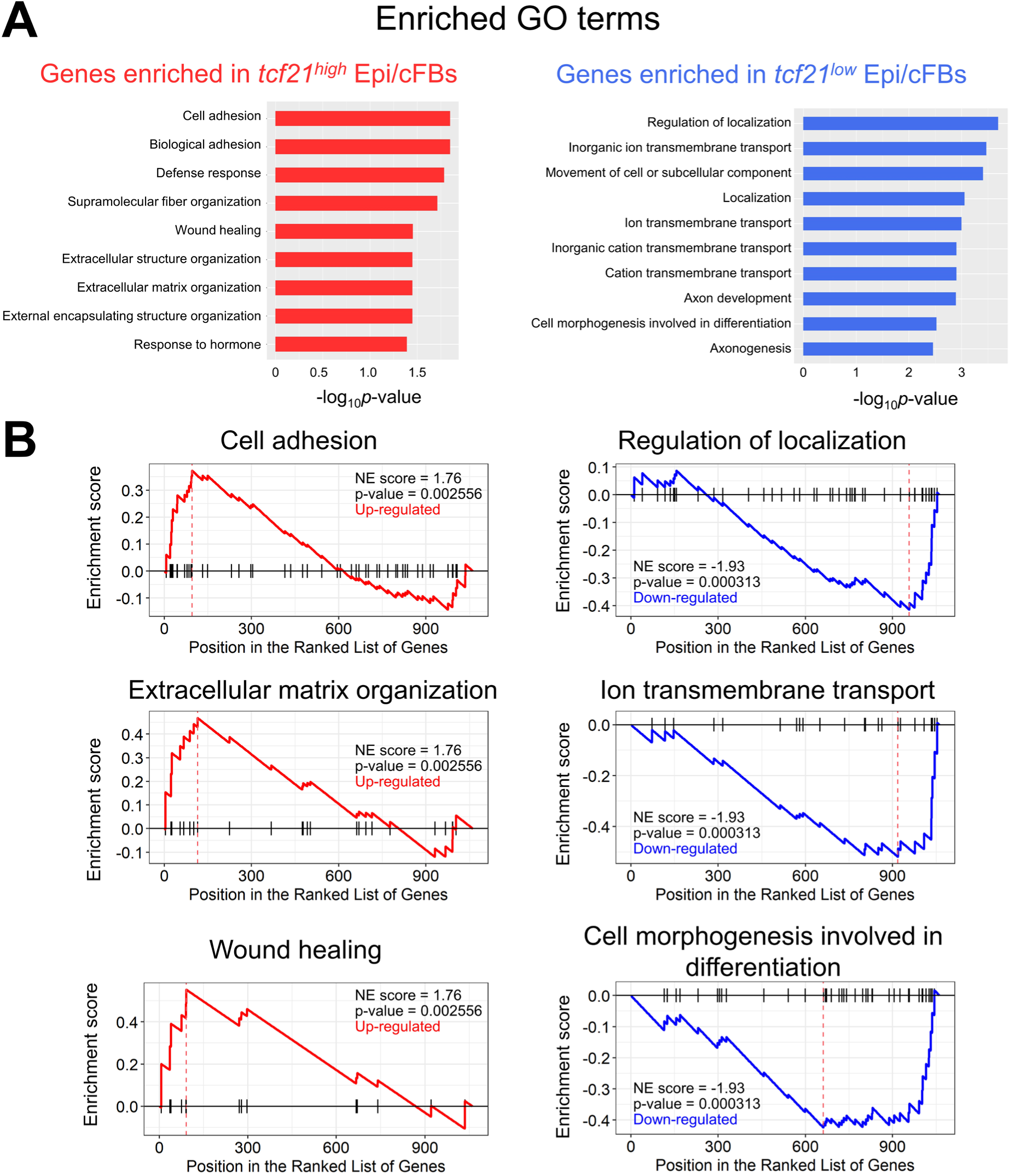
Differential gene expression analysis between *tcf21^high^* and *tcf21^low^* Epi/cFBs. (A) Top GO terms for genes enriched in *tcf21^high^* (Red) and *tcf21^low^* (Blue) activated ECs. (B) GSE analysis plots of cell adhesion, extracellular matrix organization, Wound healing, regulation of localization, ion transmembrane transport, and cell morphogenesis involved in differentiation. NE score, normalized enrichment score.

**Figure S4.**
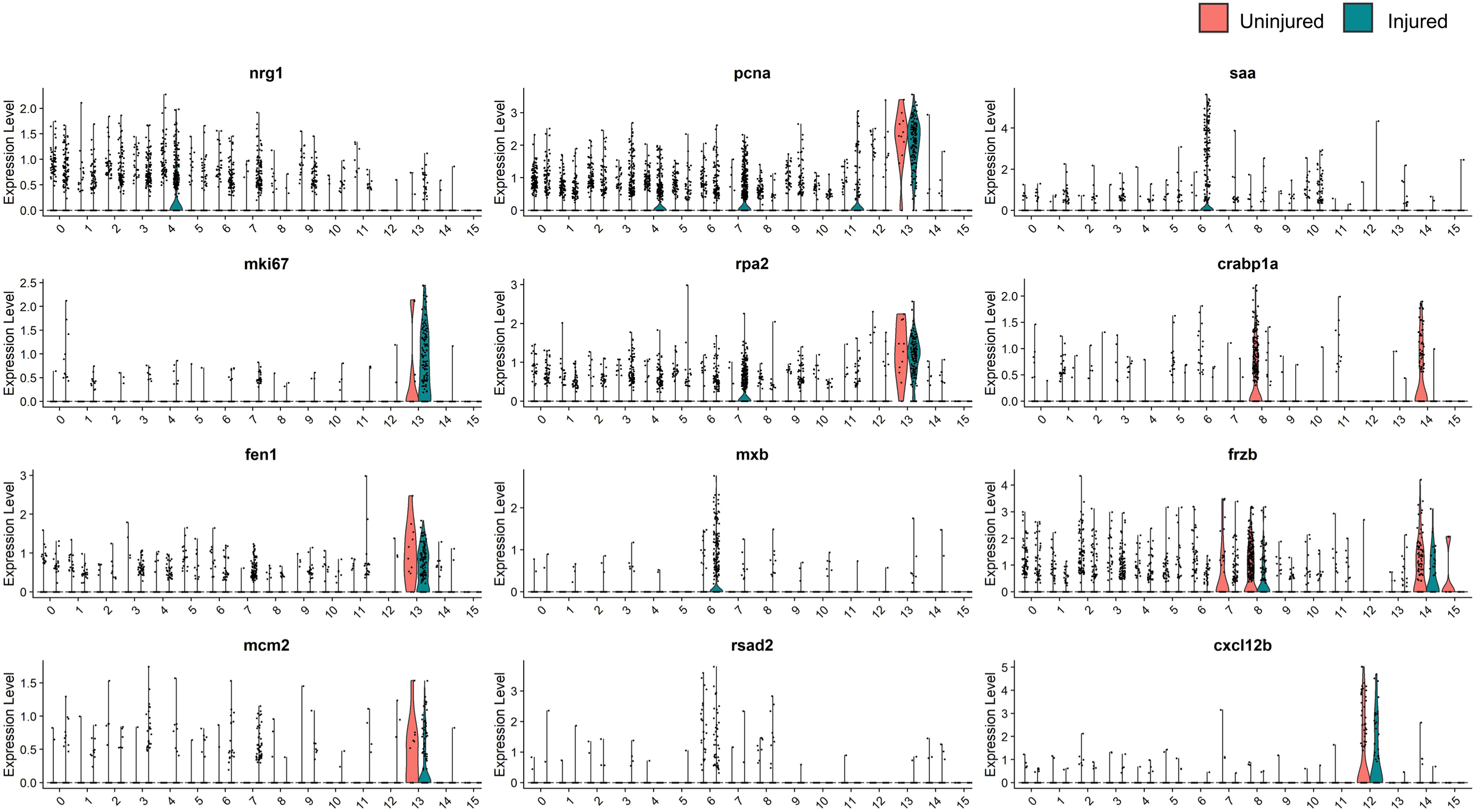
Violin plot comparing the expression of *nrg1, mki67, fen1, mcm2, PCNA, rpa2, mxb, rsad2, saa, crabp1a, Frzb,* and *cxcl12b* to label distinct subpopulations of *tcf21^+^* Epi/EPDCs.

**Figure S5.**
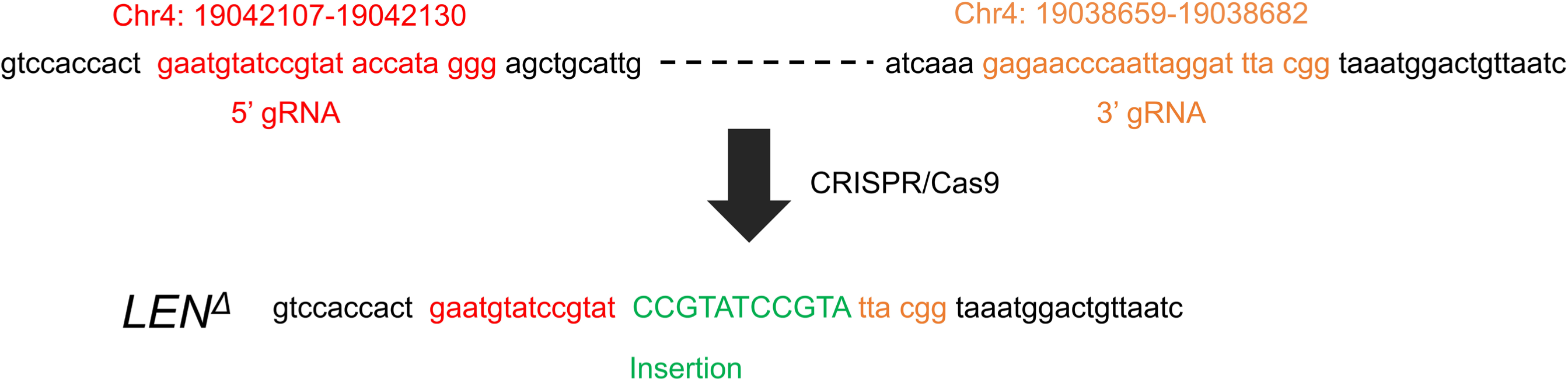
Guide RNA sequences and genomic coordinates of *LEN* deletion (*LEN^Δ^*) generated by CRISPR/Cas9. Injury-dependent expression of regenerative factors generated by *lepb^+^* activated ECs. Inj, injured. Un, uninjured.

**Figure S6.**
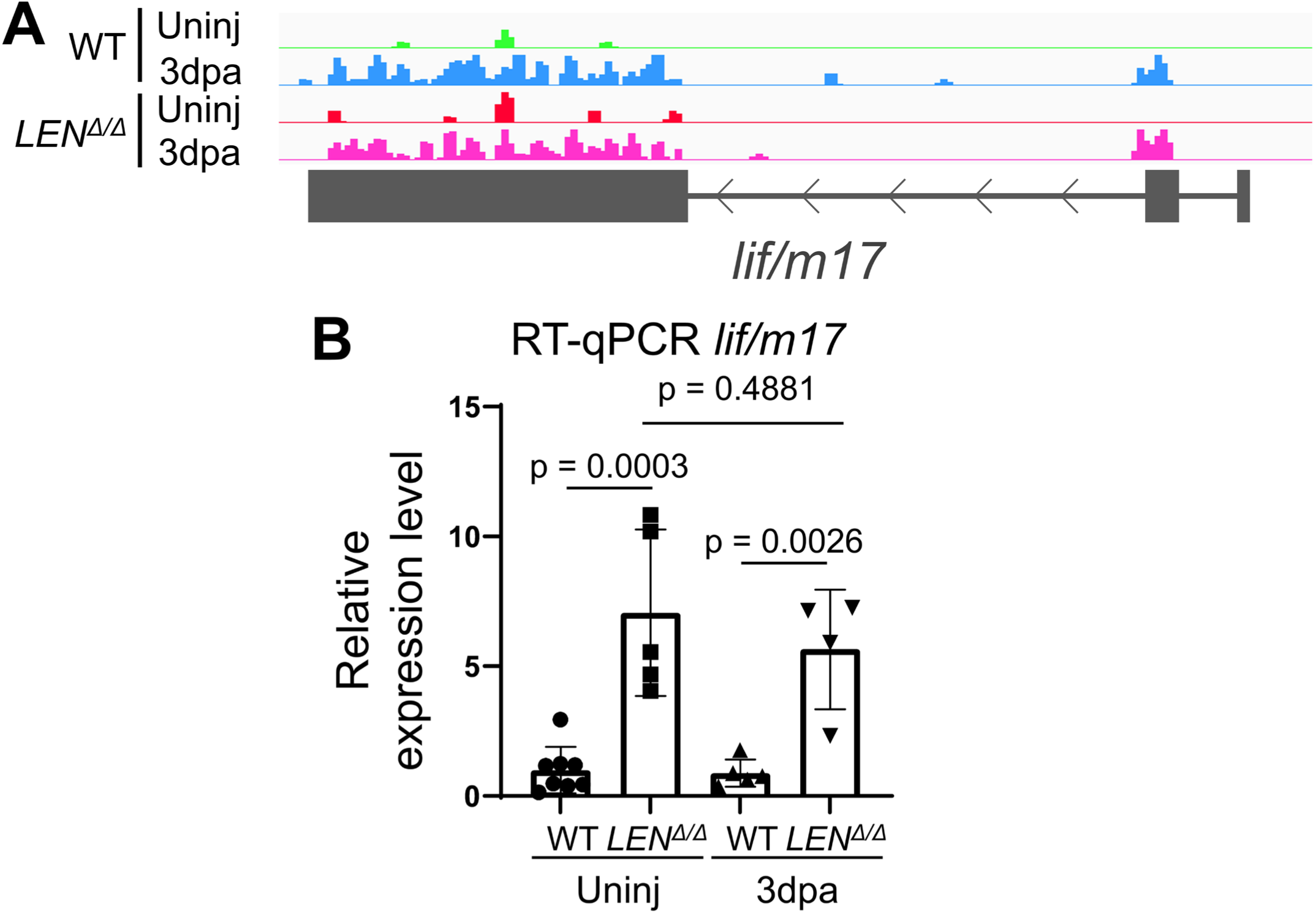
*lif/m17* is upregulated upon injury in wild-type and *LEN^Δ/Δ^* hearts. (A) RNA-seq of uninjured and 3 dpa wild-type (WT) and *LEN^Δ/Δ^* hearts. Tracks indicate *lif/m17* transcript level is increased similarly in both wild type and *LEN^Δ/Δ^* upon heart injury. (B) RT-qPCR analysis of *lif/m17*transcript level in WT and *LEN^Δ/Δ^* hearts. Data are mean ± s.d.

**Supplementary Table S1.** Cell number of *tcf21^+^* Epi/EPDC clusters in the uninjured and injured hearts identified by scRNA-seq analysis

| Cluster | Uninjured | Injured | Total |
| --- | --- | --- | --- |
| 0 | 366 | 515 | 881 |
| 1 | 370 | 437 | 807 |
| 2 | 424 | 382 | 806 |
| 3 | 601 | 201 | 802 |
| 4 | 374 | 313 | 687 |
| 5 | 266 | 412 | 678 |
| 6 | 407 | 258 | 665 |
| 7 | 526 | 40 | 566 |
| 8 | 125 | 332 | 457 |
| 9 | 252 | 190 | 442 |
| 10 | 206 | 176 | 382 |
| 11 | 126 | 167 | 293 |
| 12 | 57 | 98 | 155 |
| 13 | 138 | 11 | 149 |
| 14 | 40 | 106 | 146 |
| 15 | 10 | 8 | 18 |
| Total | 4288 | 3646 | 7934 |

## Notes

### Competing Interest Statement

The authors have declared no competing interest.

